# Development of a clot-adhesive coating to improve the performance of thrombectomy devices

**DOI:** 10.1101/2022.10.18.512663

**Authors:** Charles Skarbek, Vaia Anagnostakou, Emanuele Propocio, Mark Epshtein, Christopher M. Raskett, Romeo Romagnoli, Giorgio Iviglia, Marco Morra, Marta Antonucci, Antonino Nicoletti, Giuseppina Caligiuri, Matthew J. Gounis

## Abstract

**Background:** The first-pass complete recanalization by mechanical thrombectomy (MT) for the treatment of stroke remains limited due to the poor integration of the clot within current devices. Aspiration can help retrieval of the main clot but fails to prevent secondary embolism in the distal arterial territory. The dense meshes of extracellular DNA, recently described in stroke-related clots, might serve as an anchoring platform for MT devices.

**Objective:** Evaluate the potential of DNA reacting surface to **aid the retention of the main clot as well as of its small fragments within the thrombectomy device** and improve the potential of MT procedures.

**Methods:** Device-suitable alloy experimental samples were coated with 15 different compounds and contacted with extracellular DNA or with human peripheral whole blood, to compare their binding to DNA versus flowing blood elements, *in vitro*. Clinical-grade MT devices were coated with two selected compounds and evaluated in functional bench tests aiming to studying clot retrieval and distal emboli release, concomitant with contact aspiration, using an M1 occlusion model.

**Results:** Binding properties of samples coated with all compounds were increased for DNA (≈ 3-fold) and decreased (≈ 5-fold) for blood elements, essentially platelet, as compared to the bare alloy samples, *in vitro*. Functional testing showed that surface modification with DNA-binding compounds improved clot retrieval and significantly reduced secondary embolism during experimental recanalization of occluded artery 3D model by thrombectomy procedures.

**Conclusion:** Our results suggest that device coating with DNA-binding compounds can considerably improve the outcome of MT procedures in stroke patients.

**What is already known on this topic –** New mechanical thrombectomy device are being improved on the conformation and shape to increase the interaction clot on the physical point of view. However, none interact specifically with the structure or composition of the clot.

**What this study adds –** The design of a chemical surface modification of the device opens the way for a specific targeting tool to increase the interaction with the clot on the molecular level.

**How this study might affect research, practice or policy –** This new surface modification, which can be applied to all commercially available mechanical thrombectomy devices, leads to a decrease in secondary embolization which cannot and is not monitored during the procedure and responsible for new territory damage.

## 1 INTRODUCTION

Despite several effective preventive strategies, stroke remains the leading cause of permanent disability^1^. In the settings of acute intracranial large vessel occlusions, the current generation of mechanical thrombectomy (MT) devices has been associated with a significant clinical benefit^2, 3^. However, MT procedures carry the risk of iatrogenic clot fragmentation and embolism towards the distal vascular bed, defined as secondary embolism (SE). Such SE may unfavorably influence clinical outcome^4^. Various strategies have been employed to reduce SE rates including the design refinement of MT devices and the study of their effectiveness to interact with the clot^5–7^. The latter would be favored by devices able to selectively adhere to clot-enriched components and not to flowing blood elements. In this setting, recent studies have extensively described the presence of Neutrophil Extracellular Traps (NETs), dense meshes of extracellular DNA, consistently found around and inside the retrieved clots^8, 9^. We therefore hypothesized that engrafting of DNA-binding compound on the surface of MT device struts might improve their ability to adhere to clots and retain released clot fragments, through the binding of the associated DNA meshes.

In the present study we have screened the potential of fifteen known DNA-binding compounds in terms of specific capture of extracellular DNA versus non-specific stickiness to blood components when immobilized on device-suitable alloy discs *in vitro* and evaluated the performances of clinical-grade, surface-modified, stent-retrievers in a simulated *in vitro* middle cerebral artery (MCA) occlusion model^10^.

## 2 MATERIAL AND METHODS

### 2.1 Material

Nitinol (NiTi) flat discs (4.8 mm diameter, 0.25 mm thick) were laser cut and mirror polished by a controlled industrial workshop (Vuichard Michel SAS, Dingy-en-Vuache, France) from a flat NiTi ribbon (5 mm diameter, 0.25 mm thick, Goodfellow Cambridge Ltd, Huntingdon, UK) and were used for the *in vitro* experiments. For the bench test evaluation, eV3 solitaire devices (6 mm × 20 mm × 180 cm, Medtronic Neurovascular, Irvine, CA) were used.

### 2.2 Surface modification of NiTi material

All NiTi material were ultrasound cleaned in successive acid, alcohol, and water baths before the functionalization process comprising three successive dip-coating steps, as described in the patent application WO2021EP64257. Briefly, the discs were first immersed in an alkaline solution of dopamine (Alfa Aesar, A11136) for 20 +/− 2h under stirring, to obtain a thin polydopamine (PDA) film^11^. Deionized water washes and ultrasound sonication was applied to withdraw PDA aggregates prior to immersion in the second bath, aimed at grafting an amine functionalized-cyclooctyne derivative anchor (DBCO-Sulfo-PEG(4)-NH_2_, IRIS biotech GMH) on the free catechol group from the PDA film. After extensive washing with deionized water, the final step led to the immobilization of an azide derivative of each compound of interest, through a bio-orthogonal alkyne-azide copper free click-chemistry reaction.^12^ Uniform coating of the medical grade MT devices, eV3 solitaire devices was achieved using an automatic dip-coater (ND-DC, Nadetech, Navarra, Spain). Once coated, the medical devices were soaked in absolute ethanol for 1 min, left to dry and resheated before the experiments.

### 2.1 Surface modification characterization

Surface modification of the flat samples was characterized by X-Ray Photoelectron Spectroscopy, Atomic Force Microscopy and ζ - potential measurement, as detailed in *supplementary material*.

### 2.2 Evaluation of the binding to extracellular chromatin versus circulating blood platelets

Chromatin and platelet binding evaluation were studied as described in the patent application WO2021EP64257 ^13^ and detailed in *supplementary material*. Briefly, the amount of captured extracellular DNA or blood platelets was quantified by computer-assisted analysis of fluorescence microscopy images. The ability of coated surfaces to bind extracellular DNA was evaluated by applying the active surface of the experimental discs stained with the cell impermeant nuclear dye Sytox Green (S7020, Invitrogen, France) upon a 3-minute contact with human neutrophils stimulated with nigericin (which trigger the formation of NETs ^14^).

Immersion in fresh whole peripheral human blood for 10 minutes, followed by an immunostaining with fluorescent antibodies directed against CD61 allowed quantification of adhering blood platelets.

### 2.3 Functional bench assay - MCA occlusion model

These experiments were performed by an experienced neuro-interventional radiologist (V.A.) and aimed to evaluate the effect of surface-modified eV3 solitaire devices compared to un-modified eV3 devices (bare metal stent, BMS) in terms of clot retrieval and SE decrease. The model reproduces the conditions of a middle cerebral artery (MCA) occlusion^15^,^10^. The clot used in these experiments was prepared using thrombin-induced clotting of bovine blood and experimental clots were incubated at 37°C for 48 hours prior to use. The latter steps favored the formation of extracellular traps within the experimental clots.

Prior to initiating thrombectomy, complete vessel occlusion with TICI 0 was confirmed by fluoroscopic and direct visualization of the model. Each device was deployed at the occlusion site and remained in place for three minutes before retrieval. Clot fragments generated during MT were collected into two collection reservoirs (one for emboli to the MCA distribution and the other to the ACA distribution). The entire procedure is detailed in *supplementary material*.

Surface-modified eV3 solitaire devices included stents coated MBF and Pipe-2. BMS and PDA-coated devices were used as controls.

Ten experiments were carried out for each group (BMS, PDA, MBF and Pipe-2). The maximum number of passages (thrombectomy attempts) was limited to 3. All stents were randomized, numbering them from 1 to 40. Briefly, an AXS Catalyst 5F (Stryker, Michigan, USA) aspiration catheter connected to a Penumbra aspiration system (Alameda, California, USA) was used as an adjunctive thrombo-aspiration procedure in all experiments.

## 3 RESULTS

### 3.1 Surface modification characterization

Surface modification characterization of uncoated and functionalized NiTi disc: PDA, PDA-DBCO and PDA-DBCO-Ligand were achieved using i/ XPS to evaluate the chemical organization, and, ii/ AFM to evaluate the microscopic modification. XPS analysis are reported in **Table 1**. Regarding the BMS samples, the expected peaks of Ti, Ni, O and C were observed (data not shown). The surface chemistry of coated samples was instead completely organic, showing strong C, O, and N peaks. As expected the deposited PDA coating was laterally homogeneous and vertically thicker than XPS sampling depth, about 8 nm, in agreement with published data.^16, 17^ Its elemental percent was in accordance with published data,^16^ the same way the N/C ratio of 0.10 was also in agreement with the expected value. Coupling of DBCO yielded a slight increase of the O/C ratio reflecting the chemistry of the DBCO spacer arm, which contains polyethylene glycol (PEG) – CH_2_CH_2_O-repeating units. After ligand coupling, further slight modification of surface stoichiometry was observed and coherent with the immobilization of a new chemical and indicated successful coupling of the ligand.

**Table 1:**
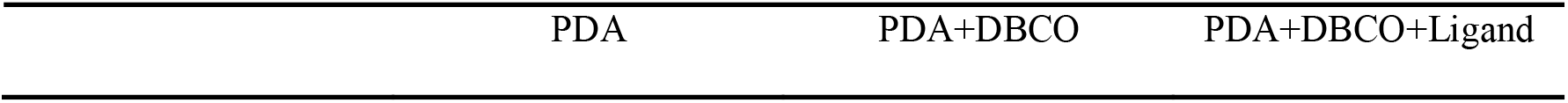
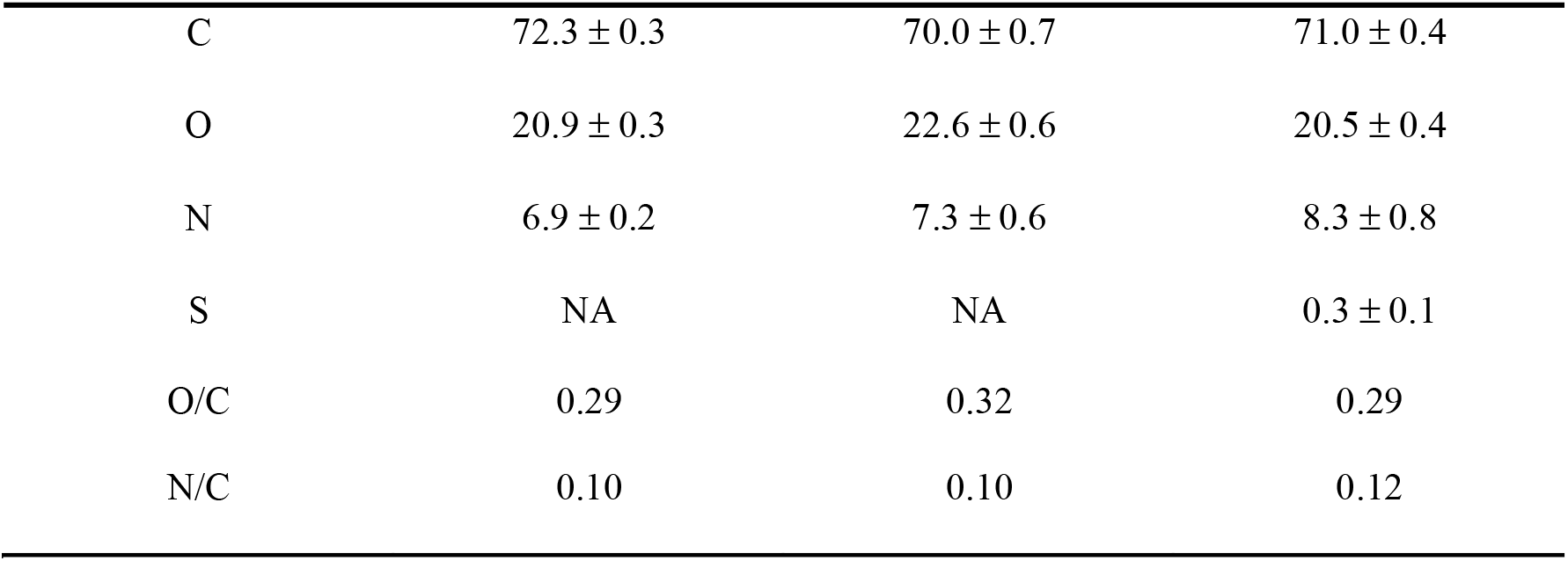
Surface organic composition of modified device-suitable alloys detected by XPS (mean ± SD; NA: Not applicable)

| PDA | PDA+DBCO | PDA+DBCO+Ligand |
| --- | --- | --- |

|  |  |  |  |
| --- | --- | --- | --- |
| C | 72.3 ± 0.3 | 70.0 ± 0.7 | 71.0 ± 0.4 |
| O | 20.9 ± 0.3 | 22.6 ± 0.6 | 20.5 ± 0.4 |
| N | 6.9 ± 0.2 | 7.3 ± 0.6 | 8.3 ± 0.8 |
| S | NA | NA | 0.3 ± 0.1 |
| O/C | 0.29 | 0.32 | 0.29 |
| N/C | 0.10 | 0.10 | 0.12 |

Noncontact AFM analysis are reported in **Supplemental information S 2**. Uncoated NiTi discs showed the presence of a flat surface with a slight difference on the morphology through AFM analysis. Regarding PDA coupling, it led to a classic spot morphology on the surface,^17, 18^ which was uniformly distributed. SEM images confirm the presence of spherical particles on PDA surface (data not shown). Adding DBCO followed by the final addition of the ligand did not modify the morphology of the surface. The measurement in three random places of final surface profile (**Supplemental information S 3**) demonstrated a symmetrical profile of the surface modification with respect to the mean line.

The contact between a solid surface and a water-based medium leads to the development of a surface charge (ζ - potential) at the interface. This charge is one of the surface characteristics which could affect the interaction between the material and the biological environment.

In particular ζ - potential was measured as a function of pH, in the 4.5 – 8.5 range, in 1 mM KCl solution (SMX). In the case of bare NiTi, the pH scan is typical of a very weak acid-base interfacial activity and it is driven by pH dependent adsorption of ions. Addition of PDA shows a negative surface which is due to the presence of more phenolic groups exposed on the surface than amine. The presence of DBCO, does not change significantly the surface charge; however, the presence of ligand makes the surface potential higher in value and much more negative it may be due to the exposition of carboxylic group which make more acidic the surface of the material **(Supplemental information S 4)**.

### 3.2 Evaluation of the chromatin and platelet binding

As the functionalization main goal was to target chromatin mesh composing acute ischemic stroke thrombi, we focused on the coating of the discs with well-known chromatin interacting compounds already used in clinics for various purposes. The binding property to chromatin and stickiness to blood elements, such as platelet, of fifteen DNA interacting agents were studied considering their toxicity, scalability and manufacturing cost (**Supplemental information S 4**). Pipe-2 (piperaquine derivative) and MBF (Mustard benzo[b]furan, a DNA mustard derivative)-coated samples showed the most interesting results in regards of both platelet and chromatin binding compared to bare NiTi discs. The binding data are summarized in **Table 2**. PDA coated discs were also evaluated to assess the binding properties of the polymer film alone. All fully coated discs demonstrated increased binding to chromatin and a low platelet adhesion compared to the bare discs (ratio 0.27) with ratio ranging from 0.67 to 6.62. The best results were obtained for the MBF compound with a high affinity to chromatin with a near 3-fold increase in chromatin binding and a near 5-fold decrease in platelet adhesion compared to the bare discs. Pipe-2 also showed interesting results with a good affinity for chromatin compared to platelet (ratio of 2.27). On the other hand, PDA coating demonstrated similar specificity for chromatin and platelet binding.

**Table 2:** Binding properties in regards to chromatin and platelets (mean +/− SEM)

| Ligand | Chromatin binding<br>(% of max) | Platelet binding<br>(% of max) | Chromatin / Platelet<br>binding ratio |
| --- | --- | --- | --- |
| Bare metal | 29.69 ± 8.63 | 90.71 ± 1.54 | 0.27 |
| PDA | 24.57 ± 8.50 | 36.32 ± 5.45 | 0.67 |
| MBF | 66.07 ± 13.21 | 9.98 ± 3.37 | 6.62 |
| Pipe-2 | 44.23 ± 10.64 | 19.45 ± 3.90 | 2.27 |

### 3.3 First Pass Recanalization and TICI Score

All devices achieved complete recanalization and TICI 3 after a maximum of three passes (**Figure 1**:). BMS and MBF coated device showed higher rates of first-pass recanalization with TICI 3 in 100% of cases, compared to PDA (7 out of 10 experiments, 70%) and Pipe-2 (9 out of 10 experiments, 90%) coated devices. Higher number of passes were required to achieve complete recanalization in the PDA group compared to the other devices, but no significant differences among devices were seen. Nevertheless, these results are to be considered with caution as the MT was fulfilled with an additional aspiration through the long sheath in the ICA. Thus, the study of clot detachment followed by its aspiration through the long sheath needs to be considered (**Supplemental information S**). The clot detachment from the device is plotted in **Figure 1**: and shows an important clot detachment in the BMS group (70% of case) compared to the other groups MBF, PDA and Pipe-2 groups in which clot detached in 20, 20 and 10% of case, respectively.

**Figure 1:**
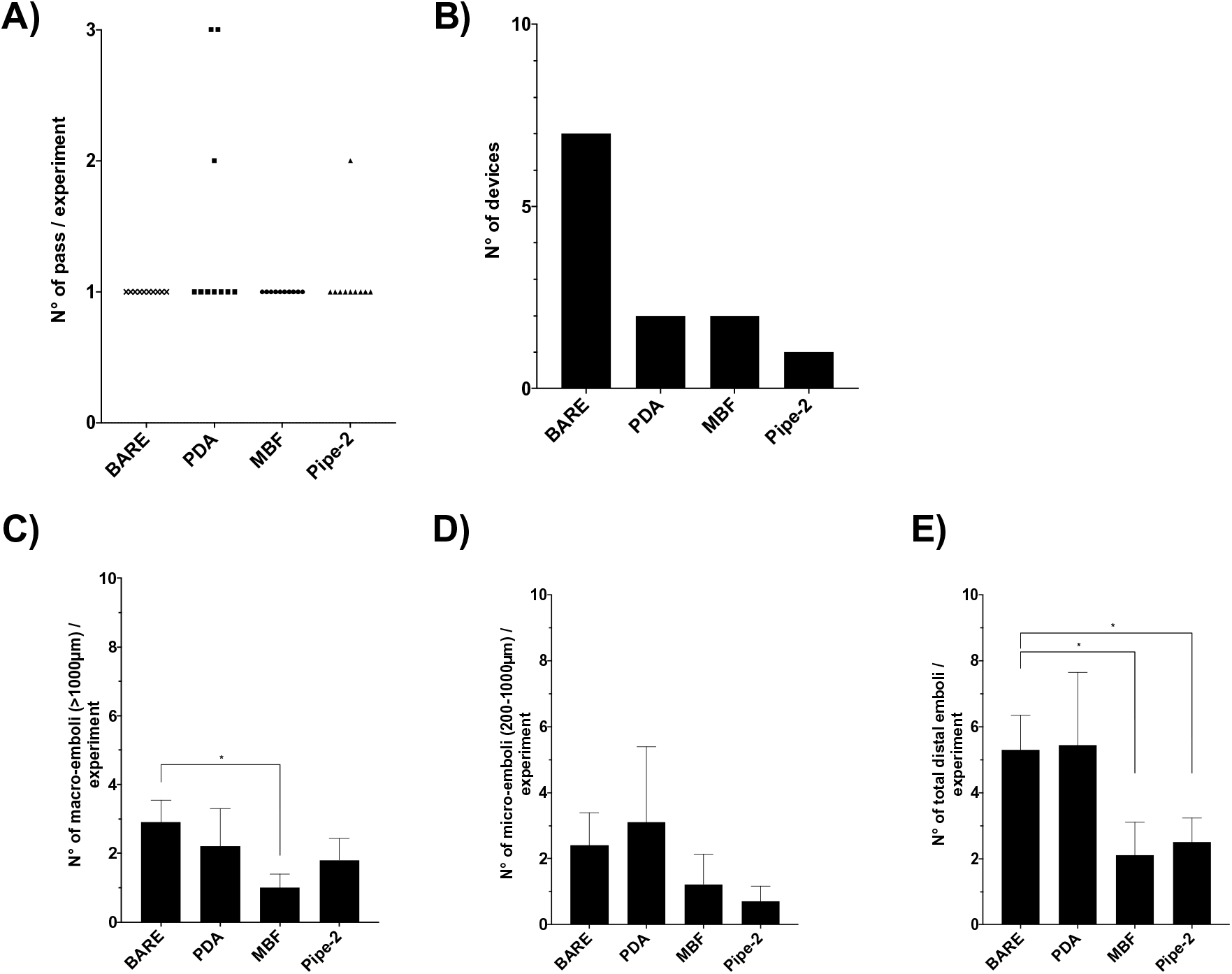
TICI score (A), clot detachment (B) and mean macro-(C) or micro-emboli (D) and total emboli (E) release during the whole study according to the tested group (n=10, * p value < 0.05).

### 3.4 Secondary Embolism

SE rates were increased in the BMS group regarding macro-, micro-emboli and overall total count of distal emboli **Table 3**.

**Table 3:** Total count of distal emboli released during the whole study (n=10) according to the tested group.

|  | BMS | PDA | MBF | Pipe-2 |
| --- | --- | --- | --- | --- |
| Overall macro-emboli | 29 | 19 | 8 | 18 |
| Overall micro-emboli | 26 | 30 | 11 | 7 |
| Overall distal emboli | 55 | 49 | 19 | 25 |

Regarding the total count of macro emboli, characterized as large clot fragments greater than 1000 μm, we found that BMS led to an increase in SE rate (29 macro-emboli) compared to the coated devices in which 19, 8, and 18 macro-emboli were recorded in the PDA, MBF, and Pipe-2 groups, respectively. The mean number of macro-emboli per experiment is illustrated in **Figure 1**:. A statistically significant difference was found for the MBF group compared to BMS (p = 0.0211).

Regarding the total count of micro-emboli, characterized as clot fragments ranging from 200 μm to 1 000 μm, BMS and PDA groups led to similar micro-emboli count with 26 and 30 micro-emboli recorded, respectively. In regards to MBF and Pipe-2 groups, a decrease in micro-emboli count compared to BMS and PDA was observed (11 and 7 micro-emboli, respectively). Regarding the mean number of macro-emboli per experiment illustrated in **Figure 1**:, no statistically significant differences were observed in either coated group compared to BMS.

Finally, regarding the overall count of distal emboli (including micro- and macro-emboli), it was found that BMS (55 distal-emboli) and PDA coated devices (49 distal-emboli) led to a higher distal emboli release compared to MBF (19 distal emboli) and Pipe-2 (25 distal emboli) coated devices. The mean number of distal emboli per experiment is illustrated in **Figure 1**:. A statistically significant difference was found for the MBF and Pipe-2 coated devices group compared to BMS (p = 0.0418 and 0.0416, respectively).

## 4 DISCUSSION

Despite the tremendous advances in the design of new MT device in the past decades,^19, 20^ MT procedure still suffers of major limitations. In spite of radiological success in about 80% of the interventions, a completely positive clinical outcome is realized only in half of the procedures.^21^ In addition, Wong et al. documented the issue regarding SE in distal or new territories which occurs in about 2 every 5 cases (40% of cases).^4^ Lastly, Luraghi et al. described the clot rolling phenomena during the MT resulting in the detachment of the thrombi from the device during the procedure, study ran with a variety of commercial BMS devices using an *in vitro* model.^22^ A recent translational study suggests that thrombectomy outcome would be significantly improved by strategies aimed at increasing the incorporation of the embolus within the device and at minimizing the release of secondary emboli.^23^ The quest for a surface modification able to selectively improve the adherence of the device to the clot matter, and not the components of the flowing blood, led us to exploit the extracellular chromatin material (only in minute amounts in the blood)that accumulates within and around occlusive thrombi,^24^ likely due to the local hemodynamic stress, consistently found in the clot retrieved from stroke patients. ^9^

Here, we report that the immobilization of DNA-binding compounds, through a scalable surface modification process applicable to any commercially available thrombectomy devices, can effectively improve the adherence of the occlusive clot to the thrombectomy device, while decreasing the release of secondary emboli, in a 3D phantom bench model of acute cerebral artery occlusion.^10^

Several compounds are known to interact with DNA, yet our *in vitro* data indicate that some of them were weaker binders of extracellular chromatin, likely according to the compound nature. The selection of the candidate for our purposes was guided the best adhesiveness shown within the relatively short contact time (3 min) and this criterion was meant to match the time allowed between the stent deployment and its retrieval during MT procedures in clinical practice. Among all the tested compounds, MBF, Pipe-2 and Pipe-4 showed *in vitro* the highest capture efficacy to DNA.

Functional tests in the bench model of cerebral artery phantom occlusion effectively showed a superiority of the modified devices in terms of clot incorporation. Indeed, the main clot readily detached from the BMS in 7 out of the 10 independent experiments whereas this occurred in a minority of the experiments performed with the coated devices (2/10, 2/10 and 1/10 with PDA, Pipe-2 and MBF devices, respectively). This observation is consistent with the study ran by Luraghi et al. describing the clot rolling phenomena during the MT when using an *in vitro* model and a variety of commercial BMS devices.^22^ Interestingly, most of the tested compounds also showed a reduced (at least a two-fold decrease) binding to blood platelets as compared to the control bare metal devices. This finding further demonstrates that the clot-capture property conferred by our surface modification is specific and supports a safer use of the modified device through the arterial bed (reduced risk of platelet aggregation onto the stent retriever). In this perspective, the mere coating with PDA, which we used as an intermediate functionalization layer, could have been proposed as a candidate for its well-known adhesive property ^25–27^. Our data however clearly show that this “sticky” property does not bring a specific binding towards DNA as compared to blood platelets: the number of total distal emboli per experiment in the PDA group was similar to the BMS, at variance with MBF and Pipe-2. This observation confirms and validates the concept that a specific binding to a component enriched in the clot, and absent in the flowing blood, can significantly improve the global performance of the thrombectomy devices.

## 5 CONCLUSION

Our work has led to the design of surface modification procedure scalable and applicable to all commercially available MT device. This work validates the hypothesis that surface modified MT device can be an interesting alternative to bare MT devices as this modification improves the capture of the main clot and to decrease the SE release.

## 6 Acknowledgments

## Contribution

CS: study design, data acquisition, data analysis, data interpretation, manuscript preparation. VA: study design, data acquisition, data analysis, data interpretation and revised the draft manuscript. AN, MJG and GC: study design, data analysis, data interpretation and revised the draft manuscript. RR: compound derivatization, synthesis, and purification. EP, ME, CMR and RM: data acquisition, data interpretation. GI, MM: data acquisition, data interpretation, manuscript preparation.

## 8 Supplementary material

### 8.1 Surface modification characterization

#### 8.1.1 X-Ray Photoelectron Spectroscopy

X-ray photoelectron spectroscopy (XPS) analysis was performed using a Perkin Elmer PHI 5600 ESCA system (PerkinElmer Inc., Waltham, Massachusetts, USA) equipped with a monochromatized Al anode operating at 10 kV and 200 W. The diameter of the analyzed spot was 500 μm; and the analyzed depth 8 nm. The base pressure was maintained at 10^−8^ Pa. The angle between the electron analyzer and the sample surface was 45°. Analysis was performed by acquiring wide-range survey spectra (0–1,000 eV binding energy) and detailed high-resolution peaks of relevant elements. Quantification of elements was accomplished using the software and sensitivity factors supplied by the manufacturer. High-resolution C1s peaks were acquired using a pass energy of 11.75 eV and a resolution of 0.100 eV/step.

#### 8.1.2 Atomic Force Microscope

Atomic force microscopy (AFM) was used to explore the surface nanotopography of 4.8 mm diameter machined Grade 4 NiTi discs untreated and treated with PDA (dip-coating step 1), PDA+DBCO (dip-coating step 2) and PDA+DBCO+Ligand (dip-coating step 3). Measurements were performed using an NX10 Park AFM instrument (Park System, Suwon, Korea), equipped with 20-bit closed-loop XY and Z flexure scanners and a noncontact cantilever PPP-NCHR 5M. This instrument implements a True Noncontact ™ mode, allowing minimization of the tip–sample interaction, resulting in tip preservation, negligible sample nanotopography modification and reduction of artifacts. On each sample, four different sample size areas were analyzed (20 × 20, 10 × 10, 5 × 5, and 1 × 1 μm) at a scan rate of 0.1 Hz.

#### 8.1.3 ζ - potential measurement

ζ-potential measurements were performed using SurPass 3, equipped with an adjustable gap cell (Anton-Paar GmbH, Graz, Austria). Measurements were conducted on 20 × 10 mm NiTi plates (NiTi foil 250 μm thickness, Sigma Aldrich), either uncoated or PDA (dip-coating step 1), PDA+DBCO (dip-coating step 2) and PDA+DBCO+ligand (dip-coating step 3) coated. Measurements were performed in a 1 mM KCl solution, according to *pH scan* method. It consists of the measurement of ζ - potential at different pH, between 4.5 and 8.5. The pH of electrolyte solution was modified automatically by the instruments using a 50 mM HCl and 50 mM NaOH. At each pH point, three measurements were performed in order to conditioning the sample, then the fourth value was taken and reported. All calculations were performed by the instrument software.

### 8.2 Evaluation of the binding to extracellular chromatin versus circulating blood platelets

Chromatin and platelet binding evaluation were studied as described in the patent application WO2021EP64257.^13^ Bare and coated NiTi discs were used within one week after their coating. The discs were washed twice with sterile PBS before running the evaluation.

#### 8.2.1 Extracellular chromatin binding evaluation

The ability of coated surfaces to bind extracellular chromatin was evaluated by applying the active surface of the experimental NiTi discs on the bottom of 48-well plates in which fresh isolated human neutrophils were stimulated with nigericin during three minutes, to trigger the formation of NETs ^14^. Briefly, neutrophils were isolated using an EasySep™ Direct Human Neutrophil Isolation Kit (Stelcell Technologies, France) from healthy donor whole blood retrieved in ACD tube. Isolated neutrophils (300 000 neutrophils, 300 μL) were incubated in 48-well plates with nigericin (30 μM, SML1779, Sigma Aldrich Chimie, France) at 37°C to promote the formation of NETs. After a 4h-incubation period, the plate was shaken 5 min at 400 rpm and centrifuged (5 mins, 500 G). Bare and coated discs were immersed in each well and left adhering to NETs for three minutes, selected to mimic the time between the stent deployment and its retrieval during MT in clinical practice. After retrieval of the disc, the discs were washed in PBS and fixed for 10 mins in 4% formalin solution. Adhering extracellular chromatin was then stained with the cell impermeant nuclear dye Sytox Green (S7020, Invitrogen, France). The NiTi discs were mounted face-down on ibidi® dishes, using ProlongGold® (Thermofisher, France) and left to dry overnight. Images were taken with an Axio Observer fluorescence microscope (Zeiss, Rueil Malmaison, France) equipped with an ORCA II Digital CCD camera (Hamamatsu Photonics, Massy, France). Fluorescence microscopy images were analyzed using Image J (National Institutes of Health) software, v1.50i for MacOS. The fluorescent intensity was normalized on the maximum observed intensity to evaluate the mean percentage of binding and compare data among the different surfaces.

#### 8.2.2 Blood platelet binding evaluation

Coated and bare NiTi discs were immersed in the wells of 48-well plates containing fresh whole peripheral human blood (200 μL) deriving from healthy donors. Blood was withdrawn in PPAK tubes (9-SCAT-II-5, cryopep, France), a thrombin inhibitor suited to accurately evaluate the reactivity of platelets *in vitro*.^28^ Contact with blood was allowed for 10 minutes, a time interval compatible with the travel of the medical device through the circulation of the patients during MT. After 10 minutes, the samples were washed and adhering platelets were revealed by fluorescence microscopy, following an immunostaining with CD61 antibody conjugated to PE (clone Y2/51, 130-124-881, Miltenyi Biotec, Auburn, CA, USA). Finally, the discs were mounted face-down on ibidi® dishes, using ProlongGold® (Thermofisher, France) and left to dry overnight. Image acquisition and fluorescence density analysis were conducted as described above.

### 8.3 Functional bench assay - MCA occlusion model

These experiments aimed to evaluate the effect of surface-modified eV3 solitaire devices compared to un-modified eV3 devices (bare metal stent, BMS) in terms of clot retrieval and SE decrease. The model reproduces the conditions of a middle cerebral artery (MCA) occlusion. Surface-modified eV3 solitaire devices included stents coated with PDA, MBF and Pipe-2. BMS were used as controls.

Ten experiments were carried out for each group (BMS, PDA, MBF and Pipe-2). The maximum number of passages (thrombectomy attempts) was limited to 3. All stents were randomized, numbering them from 1 to 40. Briefly, an AXS Catalyst 5F (Stryker, Michigan, USA) aspiration catheter connected to a Penumbra aspiration system (Alameda, California, USA) was used as an adjunctive thrombo-aspiration procedure in all experiments. The model system was composed of a human vascular replica, clot model, and a physiologically relevant mock circulation flow loop.^10^

#### 8.3.1 Vascular model

Twenty vascular replicas of the cerebral circle of Willis were built using magnetic resonance angiograms from 20 healthy patients. Informed written consent was obtained in each case. Such replicas were built using a small-batch manufacturing process previously described.^15^ Concretely, a core-shell mold was created for silicone infusion (Sylgard 184, Dow Corning, Midland MI). By dissolving the core-shell mold in xylene after silicone curing, a transparent and flexible silicone replica was obtained. To reduce the friction between the device and the silicone replica, the inner wall of the resulting replica was lubricated by coating a layer of LSR topcoat (Momentive Performance Materials, Albany NY).

#### 8.3.2 Clot formation model

A model of friable clot of medium stiffness (diameter: 4.3mm) was prepared for device testing using thrombin-induced clotting of bovine blood containing barium sulfate (1 g / 10ml blood), fibrinogen (29.7 mg in 0.7ml saline + 1.8 ml blood/barium mixture) and thrombin (50 NIHU thrombin in 0.5ml saline + CaCl_2_ added to 2.5ml blood/barium/fibrinogen mixture).This type of clot is prone to fragmentation during MT and was selected specifically to mimic the worst-case scenario with respect to clot fragmentation. The clot was then incubated at 37°C for 48 hours prior to use to increase the chance of chromatin release within the clot. The presence of extracellular traps was confirmed by DAPi and H3cit staining (**Supplemental information S 1**)

#### 8.3.3 *Ex vivo* cerebral circulation flow loop

The clot (length: ≈ 1cm) was injected into the flow loop driven by a peristaltic pump via a separate entry close to the silicone replica to form an MCA occlusion as described previously. ^29^ Briefly, an acrylic box containing the silicone replica was connected to a flow loop which comprised a peristaltic roller pump used to deliver an oscillatory flow of saline solution, with peripheral resistance provided by adjustable clamps and the flow through each segment was calibrated to mimic cerebral hemodynamics. A filter funnel (51 microns filter) was attached to the saline reservoir. A pulsatile pressure waveform similar to those observed in the human carotid arteries was generated by the pump.^30^ Flow and pressure metric were controlled using flow Sensors (Transonic Systems Inc., Ithaca, NY) and pressure transducers (Validyne Engineering, Northridge, CA) to measure the MCA and ACA artery flow and pressure. The data acquisition system was equipped with an analog-to-digital converter, and incorporated the synchronized video captured by a digital camcorder.

#### 8.3.4 Flow restoration procedure and distal emboli analysis

Prior to initiating thrombectomy, complete vessel occlusion with TICI 0 was confirmed. In the event that TICI > 0 was observed, the experiment was halted, the clot was removed, and the system was flushed before new clot placement. The experiment was excluded for analysis in cases of malposition of the clot (internal carotid artery or distal M2 segment) and if the clot was fragmented on delivery prior to MT. Co-aspiration was used during clot retrieval for all thrombectomy procedures as described above.

##### 8.3.4.1 Flow restoration procedure

All MT procedures were performed by an experienced neuro-interventional radiologist (V.A.) blinded to the device type. Briefly, a long sheath (Neuron Max 0.088 in, Penumbra, Alameda CA) was placed in the ICA and an aspiration catheter (Catalyst 5F, Stryker Neurovascular, Fremont CA) was then advanced up to the terminal ICA. Consecutively, a Rebar 18 microcatheter (Medtronic) was placed over a 0.014^..^ micro guidewire (Synchro, Stryker) and was positioned to the distal end of the occlusive clot. The guide wire was then withdrawn and replaced by the MT device. Each device was deployed at the occlusion site per the manufacturer’s instructions. Each device remained in place for three minutes before retrieval. The MT device supported by the microcatheter were then withdrawn slowly until the entrance into the aspiration catheter. This step ended when the backflow into the aspiration tubing stopped. Then the triaxial system (aspiration catheter, microcatheter, device) was locked and retrieved slowly as a whole. Additional aspiration was applied via a syringe for the duration of device pull back into the long sheath to avoid any effect from stripping the clot off the device. One device was used per experiment for a maximum of three passes. Fluoroscopic and direct visualization of the model was used to determine the modified TICI score. Angiography was not performed since the use of contrast would interfere with the particulate count.

##### 8.3.4.2 Particulate analysis for distal emboli

Prior to each experiment, a 500 ml blank specimen was collected to measure particulate matter unassociated with the thrombectomy procedure. Particle collection began immediately prior to device deployment and ended after clot removal. Clot fragments generated during MT were washed into two collection reservoirs (one for emboli to the MCA distribution and the other to the ACA distribution) for further analysis. First, particulates greater than 1,000 μm were manually separated from the collection chamber and measured with calipers. The Coulter Principle was used to characterize particulates with sizes ranging from 200 to 1,000 μm (Multisizer 4 Coulter Counter, Beckman Coulter, Inc., Brea, CA). The particle size distribution of the blank specimen was subtracted from that measured following the thrombectomy experiment.

---

**Supplemental information S 1.**
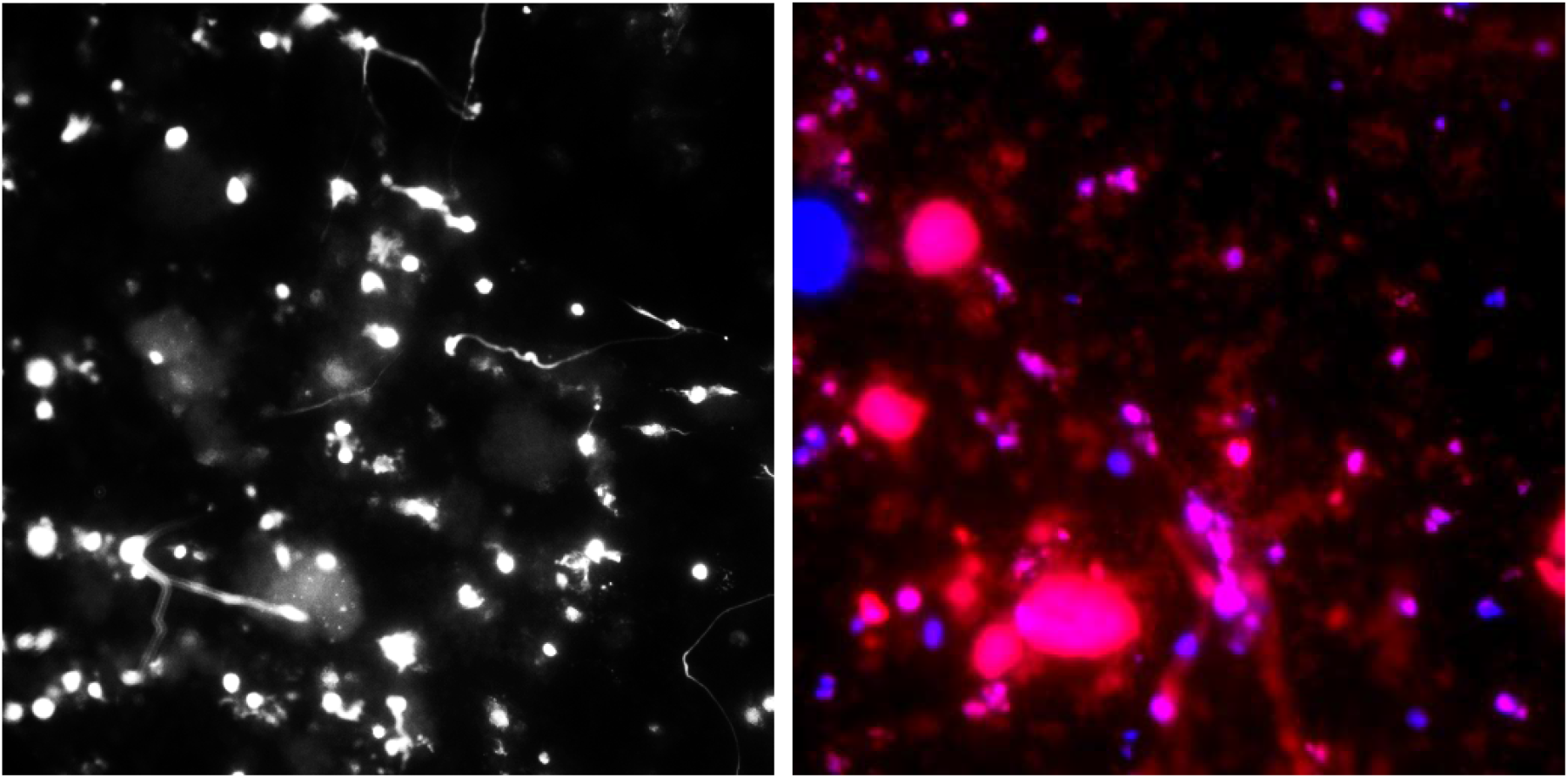
Microscopic staining clot section Left: DAPi staining alone, right DAPi and H3cit staining. Extracellular traps characterized as positive DAPi and H3cit staining confirm the presence of dense chromatin mesh within the artificial clot.

**Supplemental information S 2:**
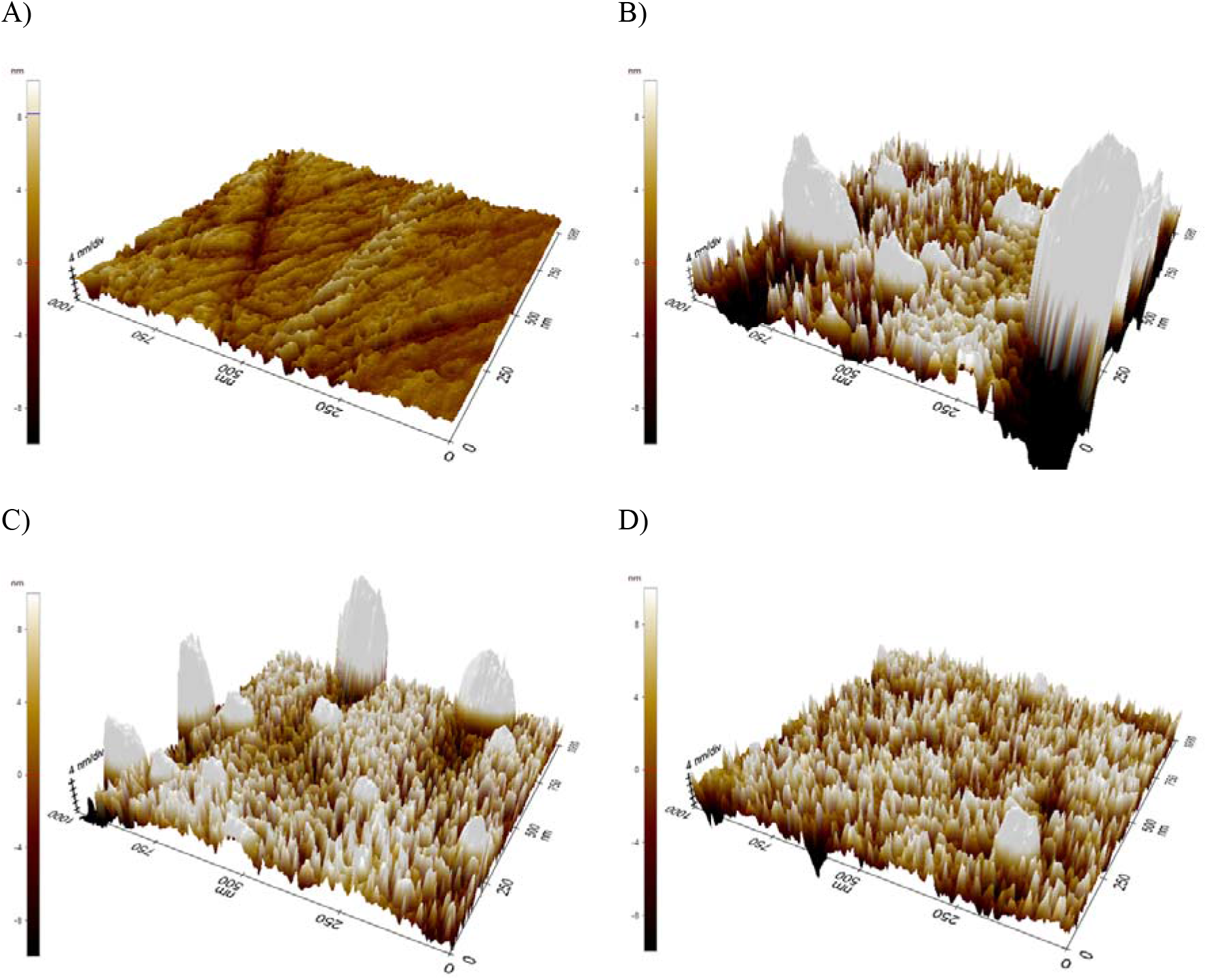
3D AFM images (1 × 1 μm) of (A) bare NiTi, (B) NiTi + PDA, (C) NiTi+PDA+DBCO, (D) NiTi+PDA+DBCO+Ligand functionalized discs.

**Supplemental information S 3:**
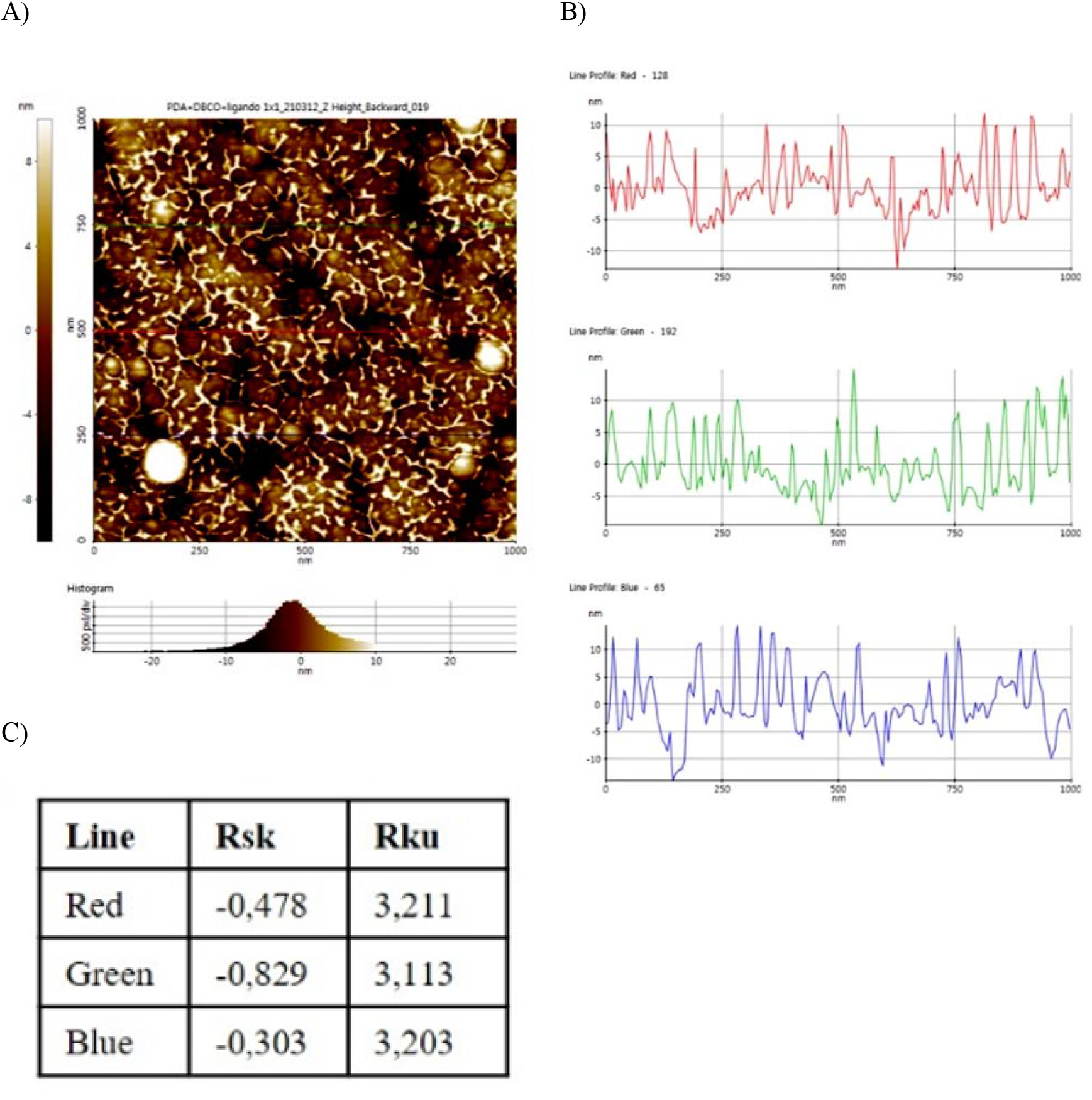
(A) AFM images of NiTi+PDA+DBCO+Ligand (1×1 μm). (B) Profile of three different sampling point. (C) Rsk and Rku value calculated on the three profiles.

**Supplemental information S 4:**
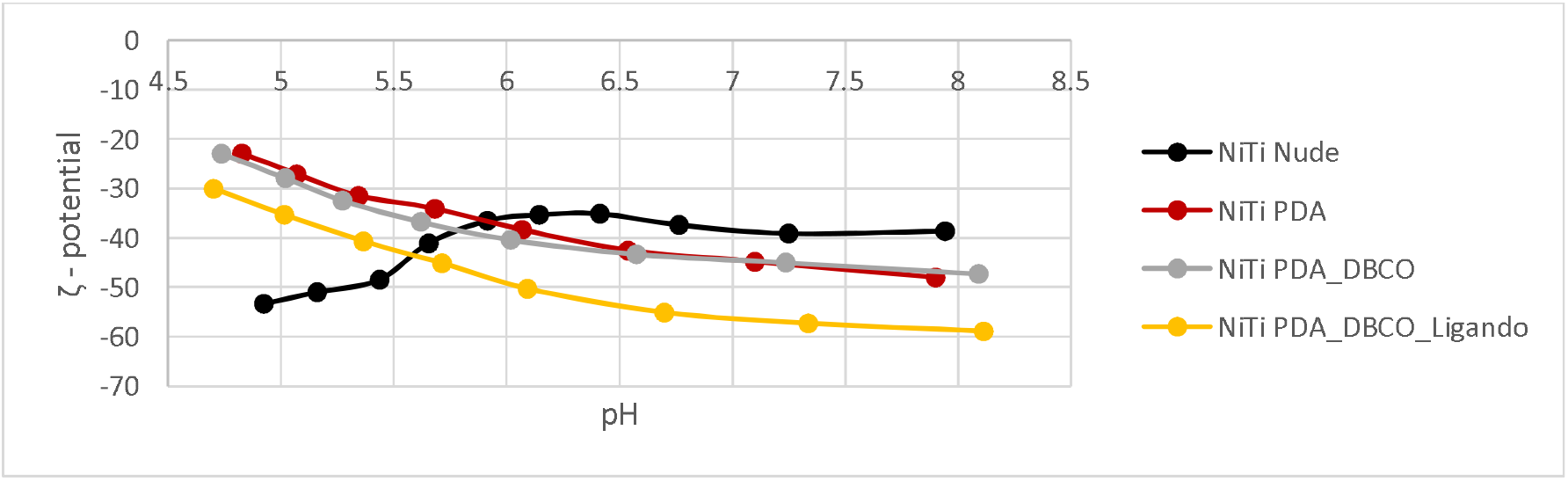
pH Scan analysis of NiTi samples, uncoated (Nude), coated with PDA, PDA-DBCO, PDA-DBCO-Ligand

**Supplemental information S 5:**
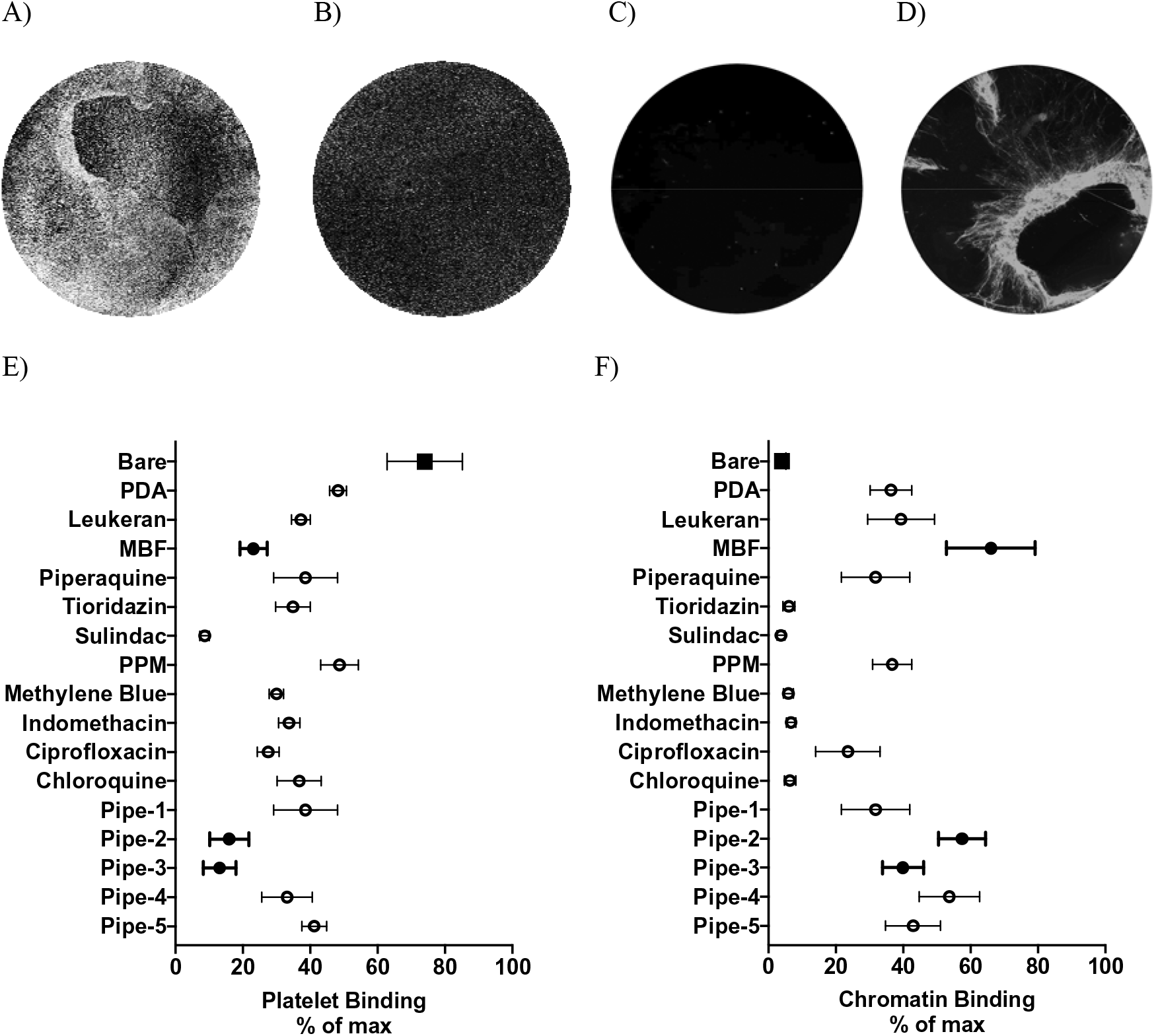
Demonstrating of the Platelet binding (A, B & E) or chromatin binding (C, D & F) on BMS (A, C) or coated (B, D) NiTi flat disc. Bare metal disc shows an increased platelet binding (A) compared to a coated disc (B). On the contrary, a coated disc (D) binds more chromatin than bare metal discs (C). E & F indicate the binding properties for each tested compound in regards of platelet binding (E) or chromatin binding (F). The best compounds are those combining a low binding to platelet as well as a high chromatin binding property (i. e. MBF and Pipe-2 coating).

**Supplemental information S 6:**
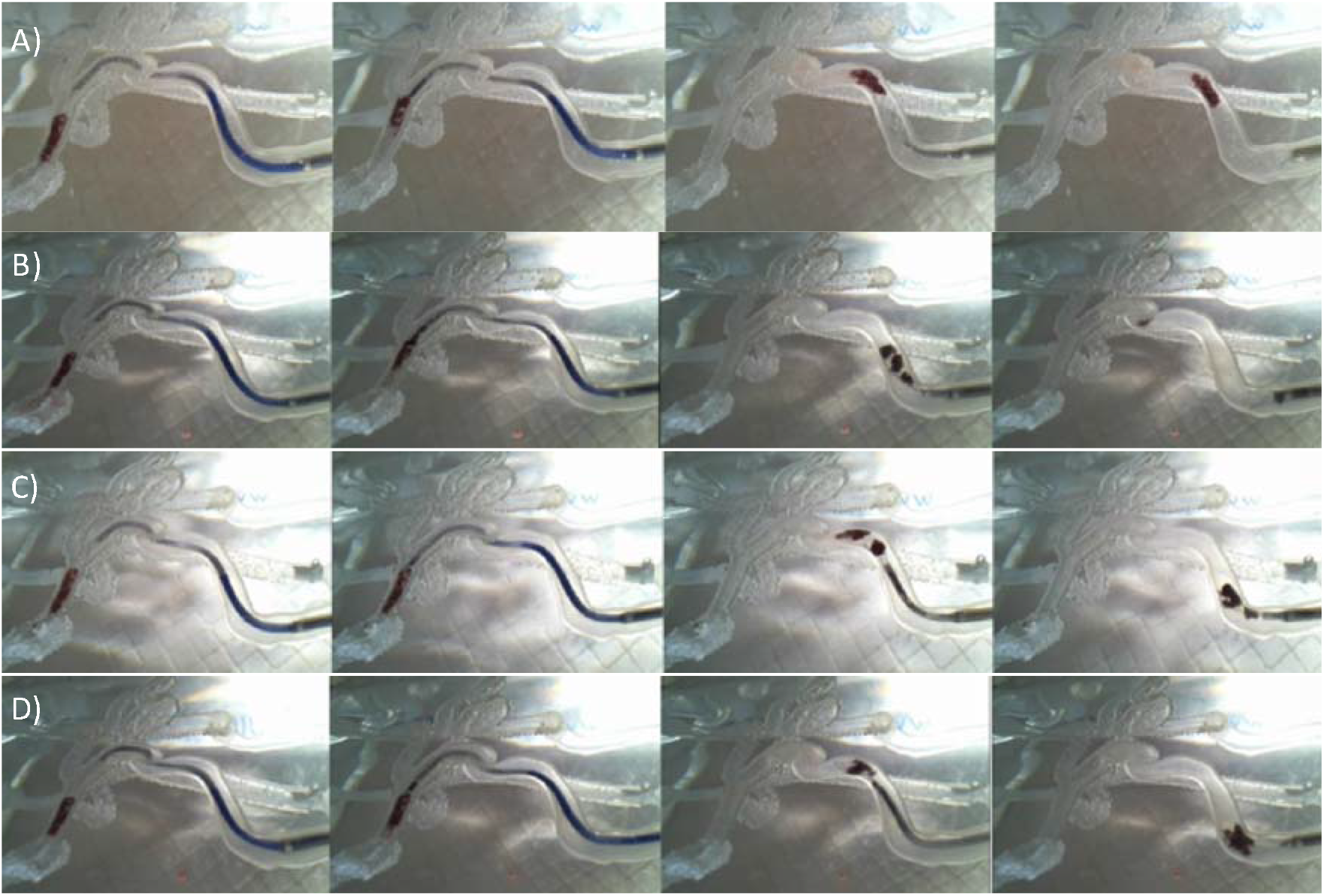
Demonstrating clot removal with the BMS (A) or coated stent with PDA (B), MBF (C) or Pipe-2 (D). From left to right and for each experiment: Stent deployment at site of occlusion (first column), followed by initiation of retrieval where the aspiration catheter engages the proximal clot surface (second column). Stent with clot during retrieval proximal to the ICA-cavernous segment (third column) and finally while entering the long sheath (fourth column). Note that the BMS complete lost contact with the clot at the end of the retrieval although the clot was aspirated into the guiding catheter giving a TICI 3 recanalization score. During retrieval with the MBF stent clot fragmentation is observed and loss of one of the fragments which is shown at the level of the siphon traveling distally. The first-pass TICI score for the specific experiment was 2. During retrieval with PDA and Pipe-2 clot fragmentation is observed with the clot fragments remaining in contact with the stent-struts until entrance into the guiding catheter.

